# Multityrosine Kinase Inhibitors Alleviate Laser-Induced Choroidal Neovascularization in Non-Human Primates: A Novel Therapy for Diabetic Retinopathy and Age-Related Macular Degeneration

**DOI:** 10.1101/2023.06.20.545744

**Authors:** Steven Dell, Ravi Nallakrishnan, Gerald Horn

## Abstract

Age-related macular degeneration (AMD) and diabetic retinopathy (DR) are leading causes of vision impairment worldwide. Both conditions involve retinal neovascularization and choroidal neovascularization (CNV), which can lead to severe vision loss. Current treatment options have limitations, necessitating the development of safer and more effective therapies. This study investigated the efficacy of Cabozantinib (CBZ), a multi-tyrosine kinase inhibitor, in a non-human primate model of retinal neovascularization. Laser-induced CNV was assessed, and CBZ demonstrated effectiveness in reducing CNV leakage and lesion area without intraocular toxicity. The inhibition of MET and VEGFR2 activation, involved in angiogenesis, is believed to be the mechanism of action. The findings support CBZ’s potential as a novel therapeutic agent for AMD and DR. Further investigations and clinical trials are warranted to evaluate CBZ’s long-term safety and efficacy in humans, as well as explore its effect on other pathways involved in CNV. The study supports the growing evidence that multi-tyrosine kinase inhibitors, including CBZ and Axitinib, hold promise in the treatment of ocular neovascularization, particularly in conditions like AMD and DR.

## Introduction

Age-related macular degeneration (AMD) and diabetic retinopathy (DR) represent two of the most prevalent causes of vision impairment and blindness worldwide. AMD primarily affects the elderly in developed countries, serving as a leading cause of blindness, while DR, a significant complication of diabetes, is a major contributor to vision loss among diabetic patients.(1,2) Central to the pathology of both AMD and DR is the phenomenon of retinal neovascularization and choroidal neovascularization (CNV). Both diseases are characterized by the abnormal growth of blood vessels in the retina, which can lead to severe vision impairment or even blindness.(3–5) The current therapeutic options for both retinal and choroidal neovascularization, encompassing strategies such as laser photocoagulation and anti-vascular endothelial growth factor (anti-VEGF) medications, possess inherent limitations. Thus, there is an imperative need for the development of safer and more efficacious treatments.. Multi-tyrosine kinase inhibitors (TKIs) like Cabozantinib (CBZ) and Axitinib have demonstrated their potency in effectively curbing angiogenesis and suppressing the activation of a variety of receptor tyrosine kinases, which includes the hepatocyte growth factor receptor (MET) and VEGFR2.(6,7) In this study, we aimed to investigate the efficacy and mechanism of CBZ in a non-human primate model of retinal neovascularization, with a focus on its potential as a novel therapeutic agent for retinal neovascularization in the context of both AMD and DR. The results of this study may provide valuable insights into the development of safer and more effective treatments for retinal neovascularization associated with these diseases.

## Methodology

### Animals

The study was conducted in a RxGen, St. Kitts Biomedical Research Foundation campus, Lower Bourryeau Estate, St. Kitts, West Indies. The study assessed the effects of CBZ on laser-induced CNV in seven adult male African green monkeys. Seven adult male African green monkeys were enrolled in the study, and baseline ophthalmic and clinical exams were performed to confirm good health and suitability for study enrollment. On day 24, after review of imaging data, monkeys were sacrificed. Laser photocoagulation was applied to six laser spots symmetrically placed within the temporal vascular arcades of both eyes, approximately 0.5-1.0 disc diameters from the fovea.(8) Animals received 4 drops (approximately 30µL per drop) of 1.5 LS CBZ malate 5mg/mL (Lot# 3-EOD-64-1/02202014@1) manufactured by Heritage Compounding Pharmacy (Brea, CA) as a ready-to-dose ophthalmic gel twice a day, two days prior to laser photocoagulation and continued for four days post-laser. Test article dosing continued once a day from day 5 to day 24.

### Ophthalmic Examination

Eyes were examined by indirect ophthalmoscopy and color fundus photography at baseline screening, immediately post-laser, and on study day 21. Fluorescence angiography and optical coherence tomography were performed on day 21.

### Criteria for Evaluation

The primary endpoints included fluorescence angiography, optical coherence tomography (OCT), and ocular biodistribution. The secondary endpoint was clinical observations.

### Fluorescence Angiogram Scoring

Late-phase fluorescein leakage at each lesion site was rated on a scale of I to IV, with IV indicating significant late leakage of fluorescein within or extending beyond the borders of the laser-induced lesion. Lesions were graded systematically from the 12 o’clock position in a clockwise manner. Scoring was conducted on each angiogram series that was enhanced in Adobe Photoshop by adjusting brightness and contrast to a similar level at each stage to minimize the impact of the inherent variability of image intensity on visual scoring as previously described.(9)

### Evaluation of Fundus Photographs

Fundus photographs were reviewed to confirm the degree of post-treatment blanching, retinal/subretinal hemorrhage and scaring, and their resolution over time.

### OCT Measurement

The CNV complex area in the OCT image was measured along the principal axis of the maximal CNV complex size within each star-shaped scan at each laser lesion.(9)

### Clinical Observations

Clinical observations were conducted at each angiogram time point to confirm the integrity of the ocular surface and normal response to mydriatics.

## RESULTS

### Topical Dosing

Six animals received bilateral topical instillations of WMD 1.5 LS CBZ malate 5mg/mL BID for 8 days (starting study day -2) then QD for 19 days starting (starting study day 6) from 2 days prior to the laser. Although no significant signs of eye irritation were observed immediately post topical dosing, a corneal cloudiness was observed in all eyes at the time of laser photocoagulation on day 0 and ophthalmic examination on day 21, which compromised the clear view for the laser treatment and fundus photography. No corneal ulcer and intraocular inflammation were observed in the study.

### Laser Photocoagulation

Six laser spots were placed in the macula of each eye. Laser photocoagulation produced acute vapor bubbles, suggestive of Bruch’s membrane rupture. Severe subretinal hemorrhage occurred in three eyes including, involving all lesion spots at the time of laser photocoagulation and not resolved by day 21; therefore, these three eyes were excluded from the study analysis. Additional 7 laser lesions, had a medium subretinal hemorrhage and were also excluded from the analysis

### Appearance of Retinal Lesions

Color fundus photography documented the appearance of retinal lesions immediately following laser photocoagulation and at study day 21. Small subretinal and retinal hemorrhage localized within or around the lesions was typically observed immediately following laser photocoagulation in the majority of eyes a component of experimental CNV induction. This small amount of hemorrhage was substantially resolved by day 21.

### Graded Angiography Scoring

Laser-induced CNV was assessed by graded scoring of day 21 fluorescence angiograms. Table 2 illustrates overall results of angiogram scoring. At day 21 no Grade IV lesions developed in a total of 35 laser lesions fitting inclusion criteria although there were 5 lesions scored as Grade IV, but excluded from the study due to medium subretinal hemorrhage. Table 2 shows comparative data in bevacizumab treated and vehicle treated control arms.

**Table 1:**
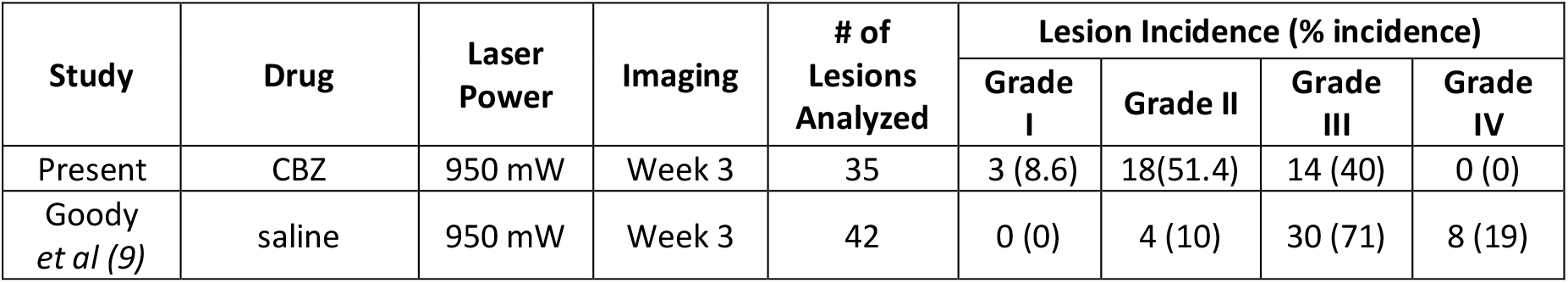
Comparison of CBZ and vehicle control.

| Study | Drug | Laser Power | Imaging | # of Lesions Analyzed | Lesion Incidence (% incidence) |  |  |  |
| --- | --- | --- | --- | --- | --- | --- | --- | --- |
|  |  |  |  |  | Grade I | Grade II | Grade III | Grade IV |
| Present | CBZ | 950 mW | Week 3 | 35 | 3 (8.6) | 18(51.4) | 14 (40) | 0 (0) |
| Goody *et al* (9) | saline | 950 mW | Week 3 | 42 | 0 (0) | 4 (10) | 30 (71) | 8 (19) |

### Quantitative Assessment of CNV Complex Area

CNV complex formation was quantified in images collected at day 21 post-laser photocoagulation with each laser photocoagulation site evaluated by area analysis of cross sectional OCT images (Figure 3). In the 35 included lesions, the mean CNV complex area was 59,391±25,503 µm^2^, ranging from 28,062 µm^2^ to 105,917 µm^2^. The mean size of CNV complex area in this study was similar to that of vehicle treated controls in previous studies employing this model.(9)

**Figure 1:**
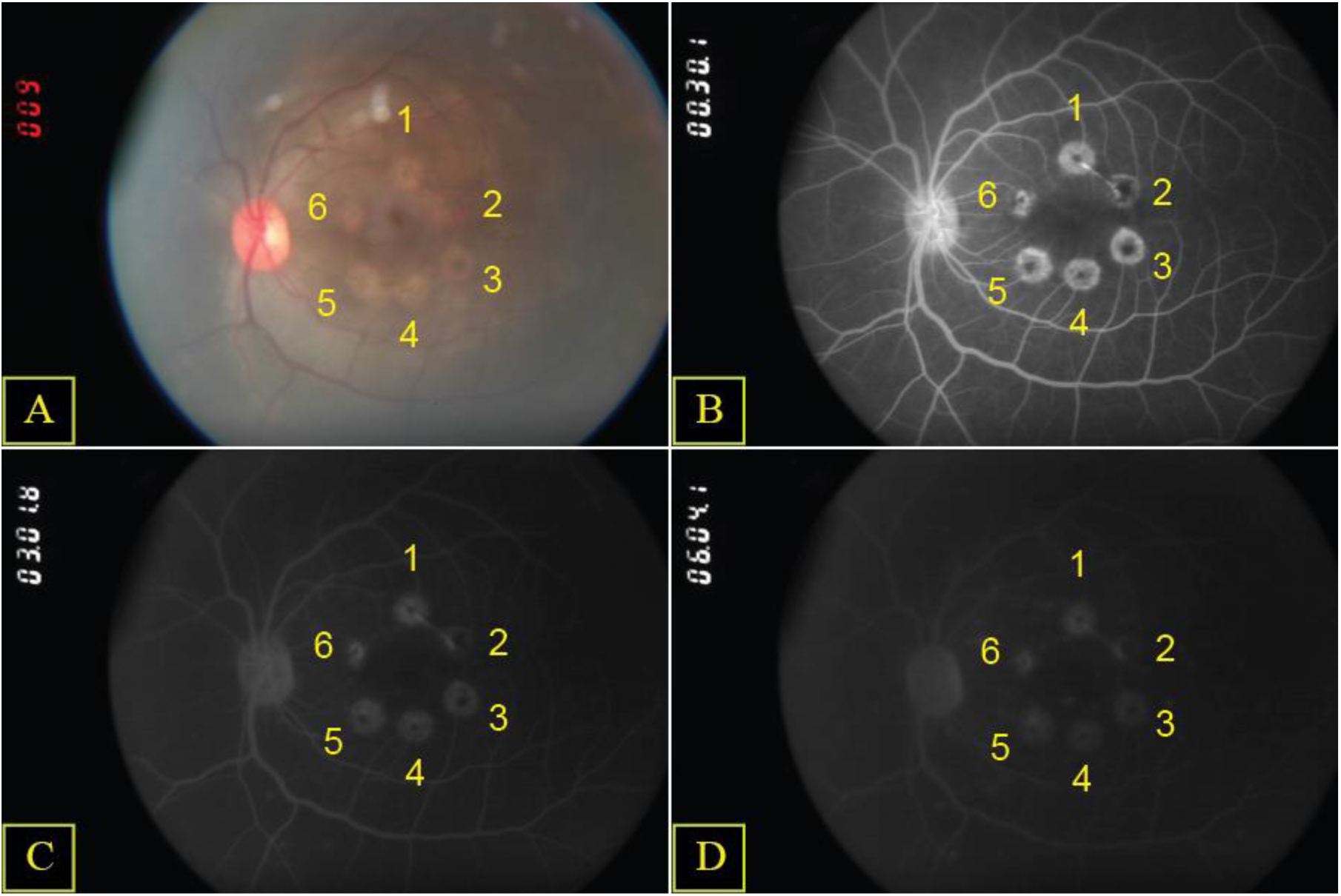
Representative CFP and FA Images at Day 21. Figure 1. Representative color fundus photograph (**A**), early phase (**B**), mid phase (**C**) and late phase (**D**) fluorescein angiograms from eye Z966 OS at day 21. Lesions were numbered systematically from the 12 o’clock position in a clockwise manner. The number illustrated in the top left corner of each angiogram is the digital time after fluorescein injection. No Grade IV lesion occurred in this eye.

**Figure 3:**
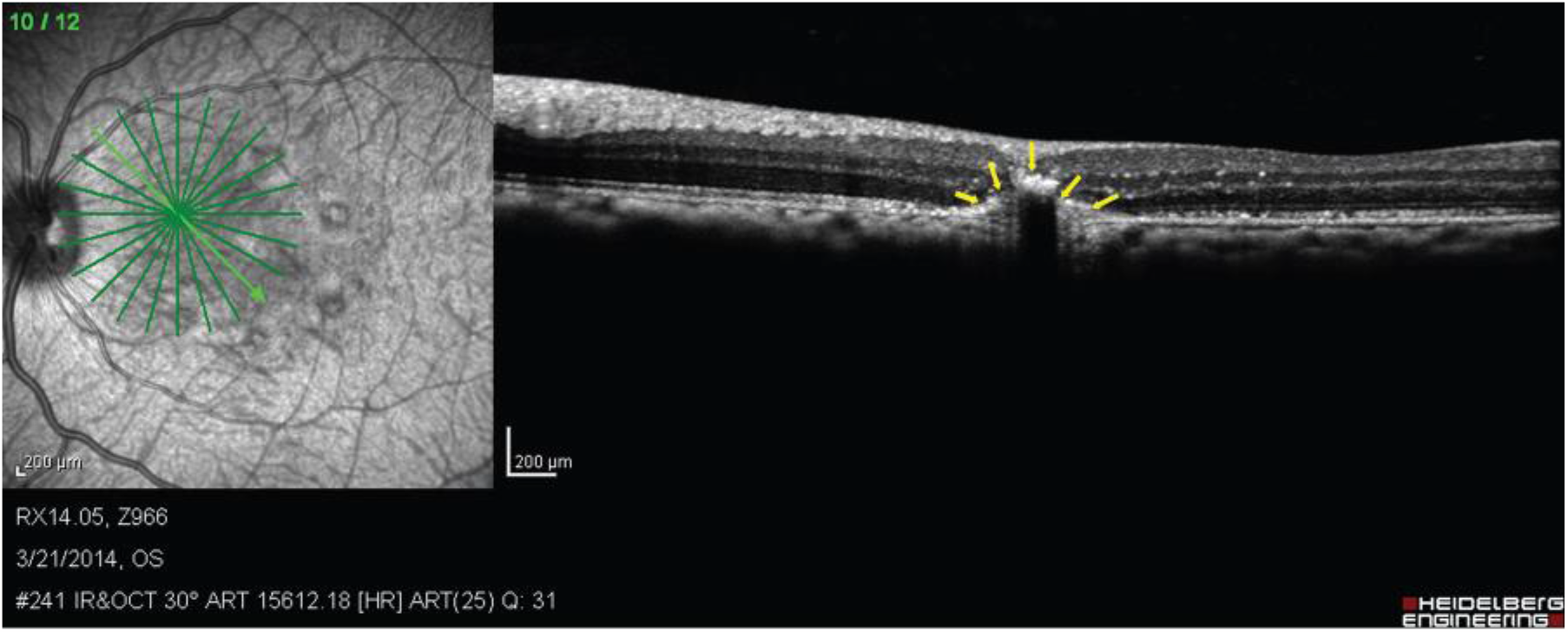
Representative OCT Images at Day 21. Figure 3. Representative OCT images. The measured CNV area is 59,723 µm^2^. Yellow arrows indicate CNV complex boundaries delineated during ImageJ analysis to calculate CNV complex area for the laser-induced lesion.

## Discussion

The current study aimed to investigate the potential of CBZ, a multi-tyrosine kinase inhibitor, in treating laser-induced choroidal neovascularization (CNV) in non-human primates. The results demonstrate that CBZ effectively reduced CNV leakage and lesion area, suggesting that it may be a novel and effective therapy for AMD and DR. AMD is a leading cause of vision loss in the elderly, and current treatment options for CNV, a common complication of AMD, have several limitations. For example, anti-VEGF drugs, which are currently the standard of care, are not effective in all patients and may have long-term side effects.(10) Therefore, there is a need for alternative therapies for CNV. The present study utilized a laser-induced CNV model in non-human primates to investigate the effects of CBZ. The results demonstrated that CBZ reduced CNV leakage and lesion area without intraocular toxicity.

The mechanism by which CBZ alleviates CNV is believed to be through the inhibition of MET and VEGFR2 activation, which are involved in tumor angiogenesis.(7) Previous studies have demonstrated the importance of these proteins in CNV, and CBZ’s ability to inhibit their activation provides a promising avenue for therapy.(5,11)

The non-human primate model utilized in the study provides a reliable and relevant model for human CNV, increasing the potential translatability of the findings to clinical trials.(12) The study’s findings provide compelling evidence for the potential of CBZ and other MTKis such as Axitinib in treating AMD and DR, Certainly, there is evidence to suggest that VEGFR-2 inhibitors, such as Pazopanib, may have therapeutic potential for diabetic retinopathy.(13) In addition to the current study, preclinical and clinical studies have demonstrated the efficacy of various TKIs, including Axitinib and Sorafenib, in reducing pathological neovascularization in models of diabetic retinopathy.(14,15) These findings suggest that TKIs targeting VEGFR-2 may represent a promising treatment option for diabetic retinopathy.

However, some limitations of the study must be acknowledged. The study was conducted on a small sample size of non-human primates, and the long-term safety and efficacy of CBZ in humans remain to be investigated. Additionally, the study did not investigate the effect of CBZ on other pathways involved in CNV, and further studies are necessary to fully elucidate the mechanism of action of CBZ in CNV.

In conclusion, our study provides preclinical evidence that Cabozantinib may be a promising therapeutic agent for the treatment of ocular neovascularization. Other MTKIs like Axitinib have also demonstrated similar potential, highlighting the importance of multitargeted kinase inhibition in ocular diseases. Future studies should continue to explore the efficacy and safety profile of Cabozantinib and other MTKIs in clinical settings to provide effective treatment options for patients with neovascular AMD and DR.

